# Disentangling cortical functional connectivity strength and topography reveals divergent roles of genes and environment

**DOI:** 10.1101/2021.04.08.438586

**Authors:** Bianca Burger, Karl-Heinz Nenning, Ernst Schwartz, Daniel S. Margulies, Alexandros Goulas, Hesheng Liu, Simon Neubauer, Justin Dauwels, Daniela Prayer, Georg Langs

## Abstract

The human brain varies across individuals in its morphology, function, and cognitive capacities. Variability is particularly high in phylogenetically modern regions associated with higher order cognitive abilities, but its relationship to the layout and strength of functional networks is poorly understood. In this study we disentangled the variability of two key aspects of functional connectivity: strength and topography. We then compared the genetic and environmental influences on these two features. Genetic contribution is heterogeneously distributed across the cortex and differs for strength and topography. In heteromodal areas genes predominantly affect the topography of networks, while their connectivity strength is shaped primarily by random environmental influence such as learning. We identified peak areas of genetic control of topography overlapping with parts of the processing stream from primary areas to network hubs in the default mode network, suggesting the coordination of spatial configurations across those processing pathways. These findings provide a detailed map of the diverse contribution of heritability and individual experience to the strength and topography of functional brain architecture.

## 1. Introduction

Evolution has shaped the cortical layout of the human brain through both scaling and reorganization ^1^. The typical globular shape of the brain evolved gradually within Homo sapiens in the last 300,000 years ^2^ and has been linked to genes associated with neurogenesis and myelination ^3^. This is related to the evolution of a developmental globularization of the brain in the first year of life, which does not occur in our closest living relatives, the chimpanzees ^4^, nor in our closest extinct relatives, the Neanderthals ^5^. It is during this developmental period that the human brain is more susceptible to environmental influence. Modern humans have more neurocranial shape variation than Neanderthals and other archaic Homo groups ^6^. A remarkable feature of the brain is that evolutionary selection and adaptation result not in a static organ, but in the capability to adapt to the environment during the life-long process of development and learning ^7^. The increased variability of functional brain architecture across individuals is associated with both differences in genetic programming and individual experience.

This variability is not evenly distributed across the brain. For instance, areas in the prefrontal cortex and association areas exhibit particularly high inter-individual differences in functional architecture ^8,9^. These areas are notable for exhibiting a predominance of long-range connectivity ^10^ as well as evolutionarily recent expansion ^8^. The link between these observations is subject to different hypotheses ^8,11^. Inter-subject variability might point at the local potential for architectural alternatives retaining comparable capacity, as it appears to be associated with plasticity and corresponding recovery of patients suffering from focal brain damage ^12^. Differences in individual experience may contribute to measurable diversity as well. Inter-subject variability in preterm neonates and healthy adults exhibit overall similar patterns ^13^, but variability increases for parts of the fronto-parietal and dorsal attention network during maturation. At the same time, genes contribute significantly to the variability of functional ^14,15^, and structural ^16,17^ cortical networks. Despite the evidence for both genetic and environmental influences on brain variability, their specific contributions to the two key features of functional organization — strength and topography — remain poorly understood.

In this study, we disentangled variability of the connectivity *strength* between components of functional networks and the spatial cortical layout of these nodes - their *topography* - to study independent contributions of genetics and environment (e.g., learning, experience) to these two features. We analysed resting state functional magnetic resonance imaging (rs-fMRI) data of twins. To disentangle strength and topography, we first performed cortical registration based on anatomical features, and then identified variability of topography by subsequent functional registration. We found that the cortical landscapes of genetic influence on these two features diverge along an axis from primary to higher order association areas. In primary areas connectivity strength exhibits high heritability compared to topography, while this relationship is reversed in association areas. Distinguishing the role of individual experience in shaping connectivity strength from the role of heritability in shaping its topography may provide crucial insights into the mechanisms enabling the flexible, yet stable nature of brain organization supporting the human cognitive repertoire.

## 2. Material & Methods

### 2.1 Dataset and preprocessing

We use the data of 231 participants labeled either as monozygotic or dizygotic twin taken from the HCP S1200 ICA-FIX denoised dataset. Only twins with 4 rsfMRI runs available are chosen. Further details on exclusion criteria and data acquisition are described elsewhere ^18–21^. 112 subjects out of 231 are labeled as monozygotic twin. The mean age in this group is 30.02 years and 74% are female. The group of monozygotic twins consists of 2 subjects denoted as “Asian/Nat. Hawaiian/Other Pacific”, 12 denoted as “Black or African Am”, 1 as “Unknown or Not Reported and 97 as “White”. The other 119 subjects labeled as dizygotic are 62% female an the mean age in this group is 29.82 years. 4 are denoted as “Asian/Nat. Hawaiian/Other Pacific”, 21 as “Black or African Am”, 1 as “Unknown or Not Reported and 93 as “White”. Participants provided verbal consent in accordance with guidelines set by the Wu-Minn HCP Consortium.

The preprocessing consists of the HCP-pipelines ^22^, which include processing of the volume data and bringing the participant’s surface into a standard space (fs_LR). For the ICA-FIX dataset independent component analysis is used to decompose the data set into “good” and “bad” components based on the volume data. Bad components are then removed from the surface data ^23,24^. Global signal regression and band-pass filtering (range 0.01 Hz-0.08 Hz) are applied in addition to the HCP-pipelines. Furthermore, the fMRI time-series are mapped to Freesurfer’s ^36^ fsaverage4 (version 5.3) surface using Connectome Workbench v1.3 ^25^.

### 2.2 Decoupling function from anatomy

In order to separate function from topography, diffusion maps are used ^26, 9^. For each twin a correlation matrix including all vertices of the left and right hemisphere is calculated. The cosine similarity metric is applied to each pair of rows of an individual’s correlation matrix and based on the obtained values, a new symmetric similarity **W** is constructed for the participant in which the entries correspond to the row pairs of the individual’s correlation matrix. Since the cosine similarity gives values between -1 and 1, the range of the entries of **W** is shifted so that the largest value is 2 and all entries are larger than 0. The matrix **W** of each individual then represents the edge weights of a connectivity graph.

The next step is obtaining a spectral representation of the connectivity structure via eigendecomposition of the matrix **L**=**Č**(**D**^-1/2^**WD**^-½^), where **D** is the diagonal matrix of node degrees d_i_=sum_j_(**W**_i,j_) and **Č** is the diagonal matrix of node degrees č_i_=sum_j_(**D**^-1/2^**WD**^-½^)_i,j_. Embedding coordinates Ψ_i_ of vertex *i* are derived from the right eigenvectors of **L** multiplied with λ_i_/(1-λ_i_), where λ_i_ denotes the eigenvalue of the i^th^ eigenvector ψ_i_, as

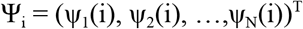

ψ_1_(i) is the i^th^ entry of the first eigenvector.

In order to reach functional correspondence across participants the embedding of each twin Ψ^S^= (ψ_1_^S^, ψ_2_^S^, …,ψ_N_^S^) is aligned to the embedding Ψ^R^= (ψ_1_^R^, ψ_2_^R^, …,ψ_N_^R^) of a reference participant (ID: 101915) not part of the twin dataset by calculating a rotation matrix **Q**_**S**,**R**_, that has the form

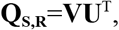

with **V** and **U** constructed via singular value decomposition,

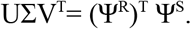

A publically available implementation is used to obtain and align diffusion maps (mapalign: https://github.com/satra/mapalign). After having obtained the embedding coordinates the first three are used as features for surface registration with MSM v2 ^27–29^, where each twin’s surface is aligned to the reference surface. Deformation fields for discovery and replicability analysis are shown in Supplementary Figure 1. After the third embedding coordinate the eigenvalues of the remaining coordinates decreased more continuously, indicating a worse separation from each other. Therefore only the first 3 coordinates were used for alignment. After alignment a vertex of the twin is assigned to each vertex of the reference participant, using nearest neighbour to reach functional correspondence of vertices across all individuals in the twin dataset.

After functional alignment, the estimation of genetic and environmental contributions to connectivity strength is repeated on the functionally aligned individuals. Additionally, the contributions to variability in spatial layout of functionally corresponding vertices are estimated. Calculations in this manuscript are performed with Python 3.6, if not stated otherwise.

### 2.3 Parcellation and connectivity matrices

After preprocessing the cortex is parceled into 600 parts, 300 per hemisphere, using the Schaefer – 600 parcellation scheme ^30^. The advantage of this scheme is the possibility to assign each region of interest (ROI) to one of the 7 Yeo-networks ^31^. In this study the networks are split into a left and a right part yielding 14 networks. For each of the 14 networks a representative ROI is chosen by correlating first each ROI time series of a network with all the other time series of the same network. The ROI with the highest mean correlation value is then chosen as the representative ROI for each of the 14 networks. For this purpose the concatenated LR-Runs 1 and 2 are used. The choice of the representative ROIs is done based on the reference subject and then taken over for all participants.

Those representative ROIs are used to describe the phenotype of the twins, which will be input of a twin model to estimate genetic and environmental influences on connectivity after anatomical and after functional alignment. Using the representative ROIs we construct connectivity matrices of size 13×13 for each vertex of the cortex by calculating the Pearson correlation coefficients between the time series of the current vertex under investigation and the 13 representative ROIs of the networks to which the current vertex does not belong. A Fisher’s z transformation is applied afterwards and gender as well as the mean relative root mean squared motion difference of Run 1 and 2 are regressed out. The regression is done using R version 3.4. To describe the phenotype of spatial layout, each functionally corresponding vertex of each participant is described by its original position in 3-dimensional space before functional alignment. Then the twin model used is the same as for connectivity strength, the only difference is that the phenotype of each participant is described by 3 traits instead of 13.

### 2.4 ACE twin model

To obtain estimates for the genetic and environmental influences on the connectivity strength of each vertex, we use a multivariate twin model ^32^. In this model the phenotype of each vertex and individual is described by 13 character traits, namely the correlation with the representative ROIs. The model is applied to each vertex separately and the relation between these character traits is summarized in the expected population covariance matrix **V**_p._ It can be split into three covariances representing the additive genetic influence **A**, the common environmental influence **C** and the random environmental influence **E**, i.e.

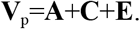

Additive genetic control refers to the additive effects of the two variants of a gene (allele) present in a human being. The two alleles can interact with each other leading to non-additive interaction effects. The typically larger additive genetic contribution together with non-additive interaction effects comprise the total genetic contribution ^33^. The common environment **C** makes members of the same family more similar to each other than to members of different families. The random environment **E** causes the differences between members of the same family. To ensure that the estimated matrices **A, C** and **E** are symmetric and positive definite, they are written as a Cholesky decomposition, **A**=**T**_A_**T**_A_^T^, **C**=**T**_C_**T**_C_^T^, **E**=**T**_E_**T**_E_^T^, with lower triangular matrices **T**_A_,**T**_C_ and **T**_E_.

To obtain estimates for **T**_A_, **T**_C_ and **T**_E_ the expected trait covariance structure of monozygotic and dizygotic twin pairs is used. It can be shown that the covariance matrices for monozygotic **V**_MZ_ and dizygotic twins **V**_DZ_ are given by the block matrices,

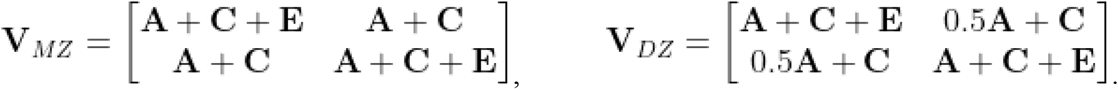

This means that the trait covariance of a twin (monozygotic, as well as dizygotic) with himself is **A**+**C**+**E=V**_p_. The trait covariance of monozygotic Twin 1 with Twin 2 is **A**+**C**, whereas for dizygotic twin pairs it is 0.5**A**+**C**, since they share on average only half of their genes, whereas the genes of monozygotic twins are identical. For parameter estimation the expected trait population covariance matrix **V**_p_, as well as the expected covariances of Twin 1 with Twin 2 (mono- and dizygotic) are then equated with the corresponding values calculated from the dataset. The twin model is implemented using R’s package OpenMx version 2.11.5 ^34, 35^.

### 2.5 Analysis

After estimation, genetic, common environmental and random environmental contribution maps are available for connectivity strength after anatomical and functional alignment and for the spatial layout. Note that values for A, C and E are only considered for vertices with corresponding fMRI signal and which can be assigned to one of the 600 ROIs (2339/2562 vertices on the left and 2341/2562 on the right hemisphere). This excludes mainly vertices from the medial wall. Several comparisons are made. First, the genetic contributions to variability in connectivity strength after anatomical and functional alignment are compared to each other by comparing the mean genetic contribution on the surface using a paired t-test. The normality assumption for the differences of paired values is verified using a histogram (Supplementary Figure 2). Additionally, the genetic contributions to connectivity strength before and after functional alignment and to spatial layout are correlated with each other using Pearson’s correlation coefficient. Furthermore, the genetic contributions to spatial layout are correlated with variability in spatial layout, distance to primary area and cortex expansion. Uncorrected and corrected confidence intervals are provided for the correlation coefficients, whereas the uncorrected confidence intervals are calculated with bootstrapping. For the t-test a corrected p-value is provided. The level of significance is set to 0.05 and obtained p-values or confidence intervals are adjusted through Bonferroni-correction. Bonferroni correction is done for each hemisphere separately.

We also performed replicability analysis using 2 RL fMRI runs, which have also been available for each participant additionally to the 2 LR runs used for the discovery analysis. In addition to performing the same steps as described in the paragraph above, we also correlated maps based on the discovery dataset with maps obtained based on the RL runs. Note that for each category either aFC, FC or SP there were some vertices for which an optimal solution for the estimates of the twin model could not be found. This holds for the replicability as well as the discovery analysis. However there were never more than 12 vertices per category for which a solution was not found. Vertices were only excluded from analyses in situations where they clearly appeared as outliers, i.e. their presence changed the appearance of the contribution maps visualized on the surface. This was only the case for maps of the variability explained by the twin model. One vertex each was removed from the discovery SP and aFC variability map as well as from the FC variability map of the replicability analysis.

### 2.6 Data and code availability

Third party code used in this study included Connectome Workbench v1.3^25^ (https://github.com/Washington-University/workbench/releases/tag/v1.3.2) and Freesurfer v5.3^36^ (https://surfer.nmr.mgh.harvard.edu/fswiki/DownloadAndInstall5.3) for preprocessing and MSM v2^27–29^ (https://www.doc.ic.ac.uk/~ecr05/MSM_HOCR_v2/) and mapalign (https://github.com/satra/mapalign) for functional alignment. Own code was developed in R 3.4 for setting up the twin model using R’s package OpenMx version 2.11.5 ^34, 35^ (https://openmx.ssri.psu.edu/), as well as in Python 3.6 for calculation of connectivity matrices. Matlab 2018 was used for generation of figures. Code and data generated during this study will be made publicly available after acceptance.

## 3. Results

To independently investigate the genetic- and environmental contribution to functional connectivity and its spatial topography across the cortex, we first disentangled these two components (Figure 1a). After anatomical alignment of cortical surfaces of all individuals ^22^, we embedded the cortical functional connectivity structure into a representational space, and aligned functional networks in this space ^37,38^. Based on these matched representations we performed diffeomorphic registration of the cortical surfaces to align regions with similar connectivity profiles^9^. This functional alignment allowed us to observe variability in the spatial position - *topography* - of corresponding functional regions (SP) and variability of functional connectivity strength of these corresponding functional regions (FC) independently (Figure 1b). We then analysed additive genetic (A), common environmental (C), and random environmental (E) factors on these two types of variability. To enable comparison of results with prior work, we also analysed these factors for functional connectivity before disentanglement (aFC). This reflects the combined variability of topography and connectivity strength at any cortical position.

**Figure 1:**
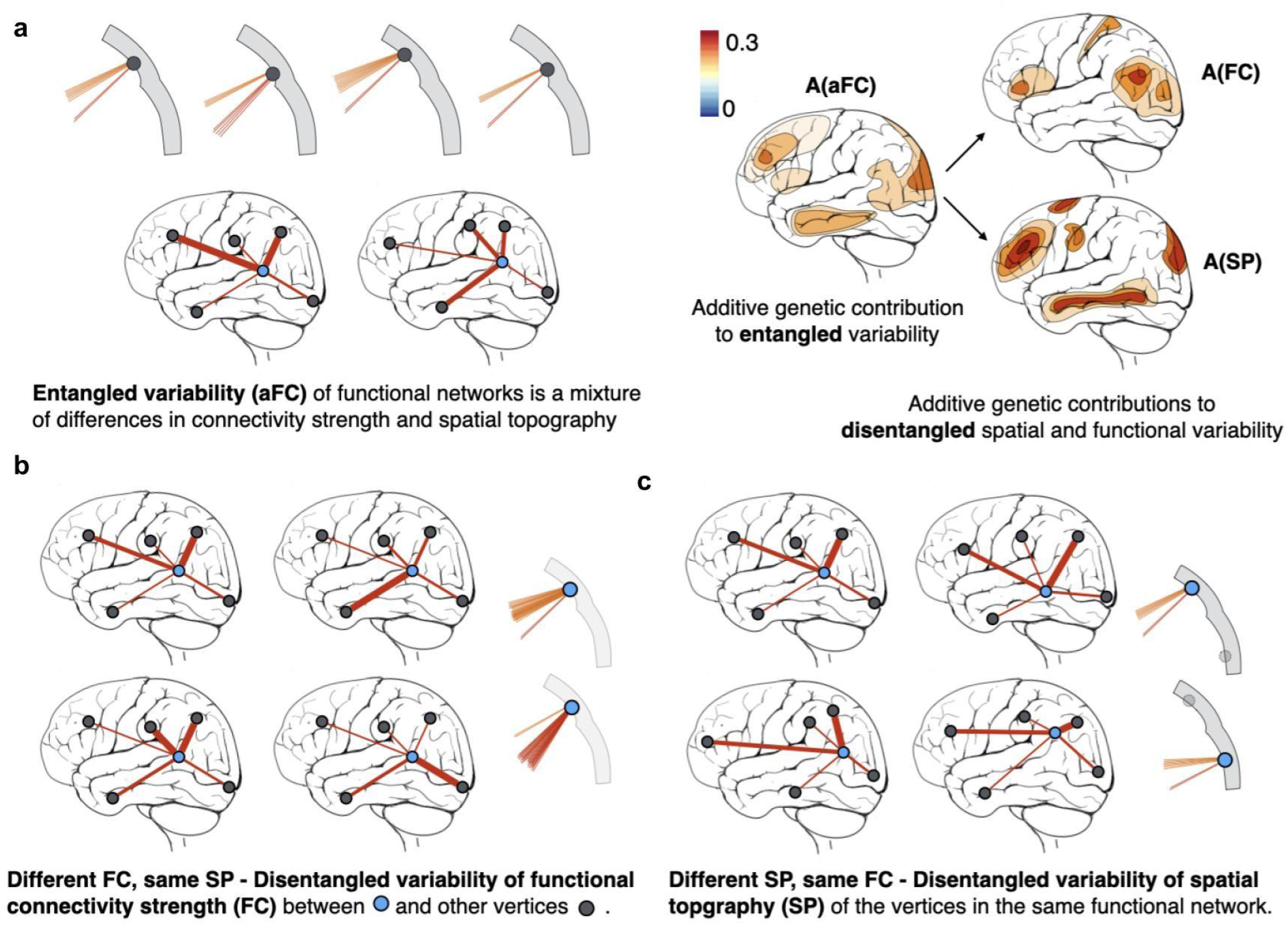
Overview. (a) Disentangling functional connectivity and topography of functionally corresponding units on the cortex enables the independent analysis of genetic contributions to these different features of variability. (b) After disentanglement, FC captures variability of connectivity strength of corresponding network nodes regardless of their spatial variability. In each of the four exemplary brains, the strength of connections is represented by the thickness of connecting lines. (c) SP is the variability of the spatial position (topography) of functionally corresponding vertices. When subjects are anatomically aligned, functional corresponding units or vertices (the blue vertex in the bottom right panel) differ in spatial position (coordinates). To know which units are functionally corresponding across subjects we perform functional alignment.

### 3.1 Divergent roles of genes and random environment in shaping strength and topography of networks

Genetic and common environmental contributions to aFC were consistent with previous findings ^15^. For aFC, areas with high non-random contributions include the rostral middle frontal cortex, the pericalcarine cortex and the boundaries between temporal middle frontal inferior parietal cortex and superior temporal banks on both hemispheres, as well as some spots in the superior frontal cortex. The transverse temporal cortex, isthmus and several region boundary areas on the medial surface exhibit highest values of random contribution (Figure 2). Surface maps for both hemispheres are provided in Supplementary Figure 3. Genetic contribution ranges from 5.25% to 45.3% of the variance explained by the twin model.

**Figure 2:**
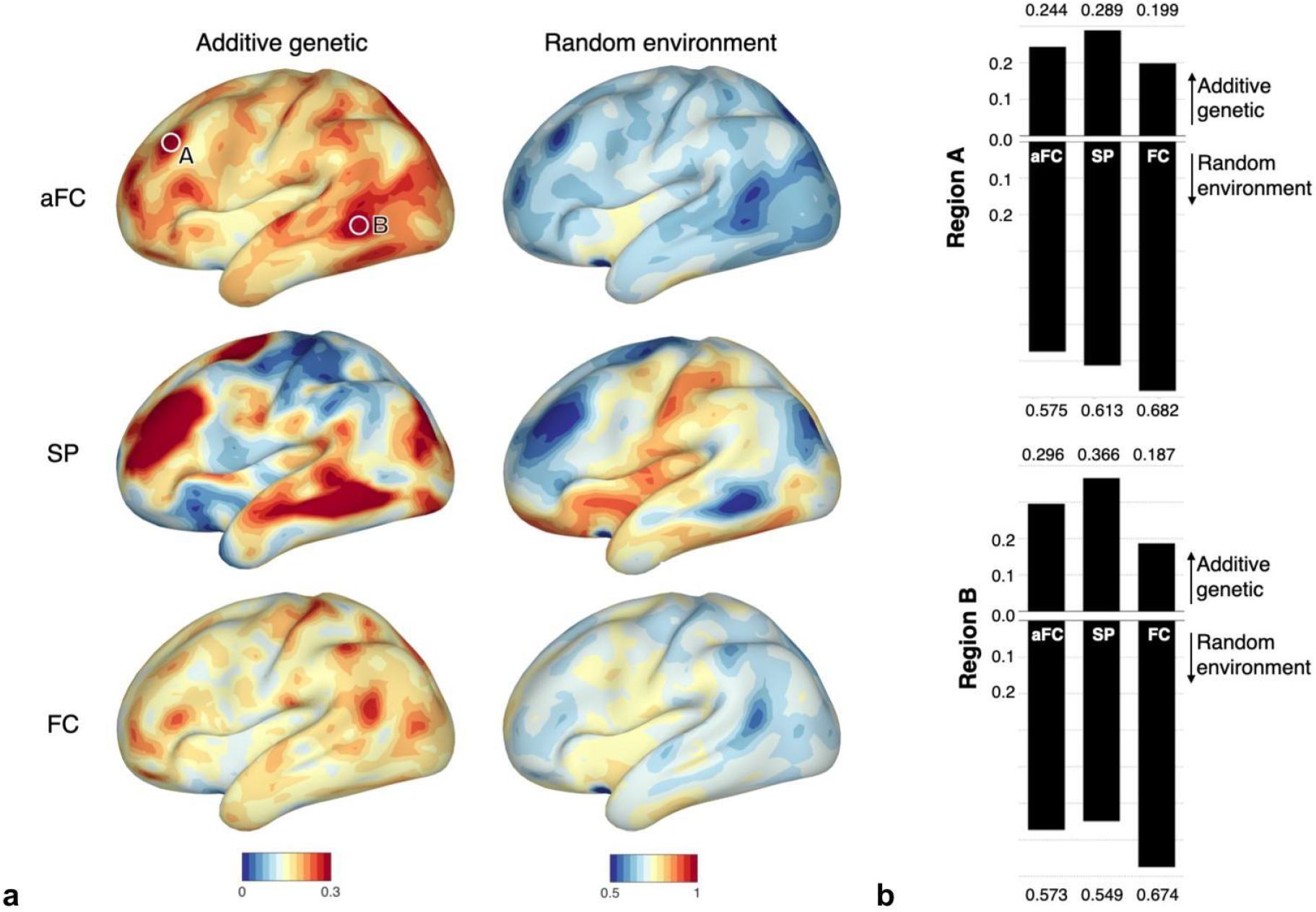
Additive genetic and random environmental contribution to functional variability of the cortex. (a) After disentangling variability of spatial topography (SP) and connectivity strength (FC), SP exhibits pronounced peaks of genetic contribution of at least 30%, whereas the genetic contribution to FC mainly decreases compared to entangled functional variability (aFC). (b) For two regions of interest indicated in the aFC map, quantitative values are shown. Compared to aFC, genetic contribution is lower for FC and higher for SP. The contribution of random environment is highest for disentangled connectivity strength FC.

After disentangling variability of function and spatial topography, genetic contributions to SP and FC diverge, and reveal a heterogeneous landscape across the cortex. In Figure 2 two example regions at peaks of A_aFC_ illustrate this divergence. On average, genetic contribution to FC is lower compared to aFC (left hem: 19.3% (A_aFC_) vs 16.5 % (A_FC_) of variance explained by genes, corrected p-value < 0.0001; right hem: 19% (A_aFC_) vs 16.1% (A_FC_) of variance explained by genes, corrected p-value < 0.0001, Figure 3a). This decrease of genetic control is also apparent in Regions A and B in Figure 2. The central sulcus is one of the few areas where local estimates of genetic contribution increased from A_aFC_ to A_FC_. Random environment is the strongest influence on FC throughout the cortex with a lowest value of 45%, while genetic contribution ranges from 2.92% to 42%. In contrast to that, A_SP_ exhibits pronounced regional heterogeneity with genetic contribution accounting from 0% to 67% of the variance explained by the twin model, giving a mean genetic contribution of 15.53% of the variance explained on the left and 14.78% on the right hemisphere. In Region A and B in Figure 2 the decrease from A_aFC_ to A_FC_ is mirrored by an increase from A_aFC_ to A_SP_.

**Figure 3:**
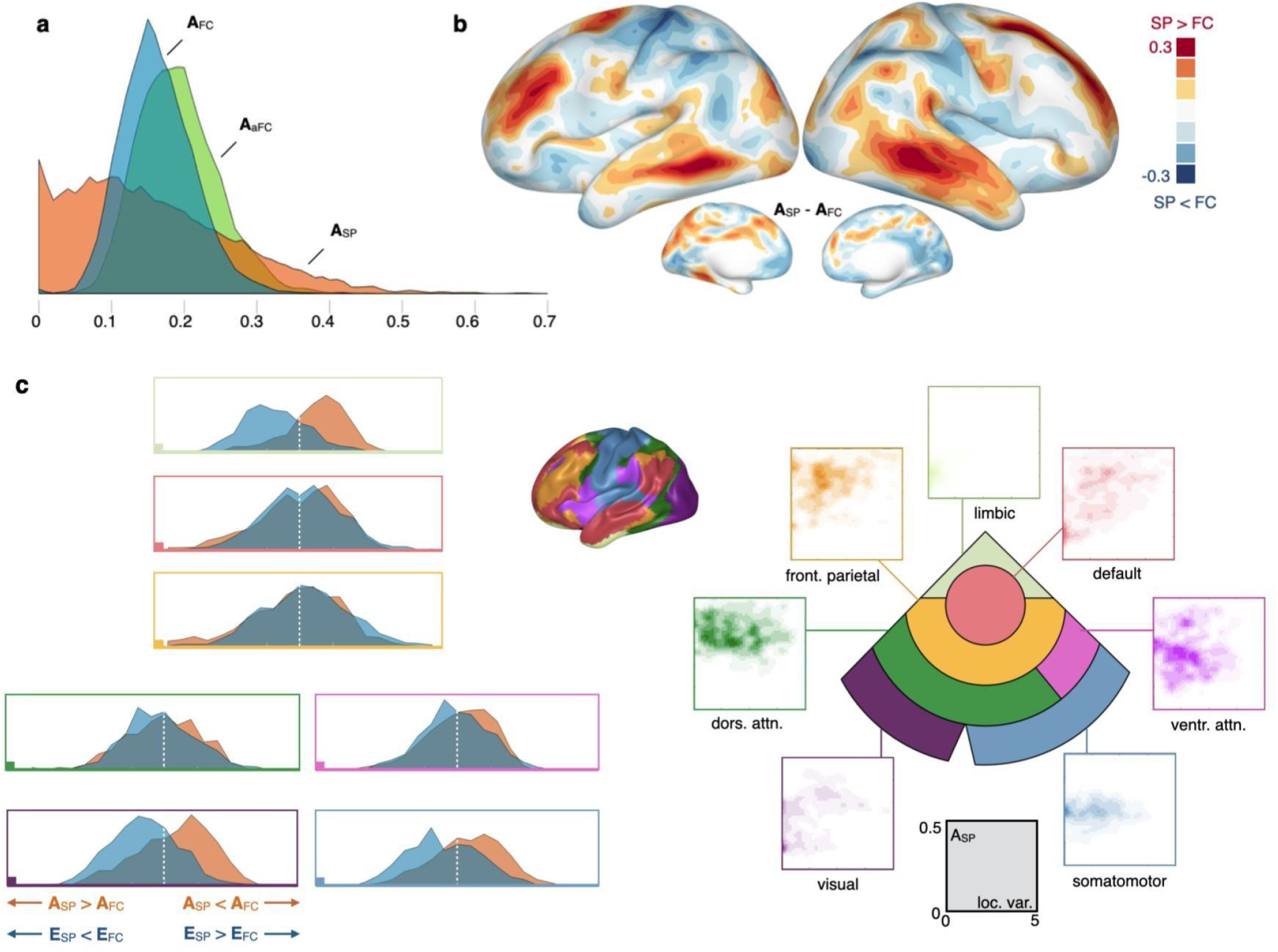
Divergent roles of heritability for disentangled connectivity strength (FC) and spatial topography (SP) of function along a gradient from primary- to heteromodal areas. (a) Global distribution of additive genetic influence A for aFC, FC and SP. Compared to aFC, genetic contribution is smaller for FC, and exhibits a more heterogeneous landscape for SP. (b) Comparing genetic contribution to FC and SP across the cortex reveals areas of dominant influence on FC (visual- and somatomotor cortex) and areas of dominant influence on SP (attention-, fronto-parietal-, and default mode network) in 7 networks (Yeo et al. 2011). (c) Heritability of connectivity strength is most dominant in primary areas (visual, somatomotor) decreasing to a minimum in integration areas (frontoparietal, default mode). On the right, for each network, the distribution of location variability and genetic contribution is plotted.

### 3.2 Two opposing gradients of genetic- and environmental influence on network strength and topography

We observed two opposing gradients in the genetic contribution to connectivity strength (A_FC_) and topography (A_SP_) along an axis from primary areas to heteromodal networks. Figure 3b shows the difference between A_SP_ and A_FC_ across the cortex. In primary areas, including visual- and somatomotor networks genetic contribution to functional connectivity strength A_FC_ dominates, while its contribution to topography A_SP_ is comparably low. The opposite is the case in higher-order areas, such as in frontoparietal areas, attention- and default mode networks ^31^ where genetic contribution to topography A_SP_ is higher than A_FC_. This coincides with a particularly high divergence between A_SP_ and A_FC,_ and A_SP_ being at its peaks higher than E_SP_ (Supplementary Figure 4). Those areas include parts of the rostral middle frontal, middle temporal and inferior parietal cortex on the left hemisphere, as well as at the boundary between the superior frontal and the precentral cortex. On the right hemisphere those areas include a part of the middle temporal cortex and an area along the boundary between superior frontal and caudal middle frontal cortex, reaching in the rostral middle frontal cortex. For 7 Yeo networks, visual- and somatomotor cortex exhibit predominantly areas with high A_FC_ / low A_SP_ paired with low E_FC_ / high E_SP_ (Figure 3c). This divergence is reduced in dorsal- and ventral attention networks, and vanishes in fronto-parietal- and default mode networks. A replication experiment shows that this observation is highly stable (Supplementary Figure 5).

### 3.3 High genetic contribution to topography in more variable, and phylogenetically modern areas

We investigated the relationship of A_SP_ to network position variability ^9^, the distance to primary areas and the amount of cortical expansion between macaque and human ^39^. A_SP_ correlates with variability (left hemi: r = 0.3, corrected confidence interval CI=(0.24,0.35); right hemi: r=0.41, corrected CI=(0.36,0.46)) and cortical expansion from macaque to human (left: r = 0.18, corrected CI=(0.11,0.24); right: r=0.099, corrected CI=(0.03,0.15)). The relationship between A_SP_ and the distance to primary area follows an inverted U shape. It increases with distance to primary areas in somatomotor-, somatosensory-, visual cortex, dorsal and lateral attention network (left: r = 0.206, corrected CI=(0.13,0.27); right: r=0.286, corrected CI=(0.21,0.35)), then decreases in limbic, fronto-parietal and default mode network (left: r = -0.275, corrected CI=(−0.35,-0.22); right: r=-0.01, corrected CI=(−0.09, 0.06)). This decrease is only significant for the left hemisphere. Supplementary Table 1 summarizes the values, and confidence intervals. Figure 4a shows the dominant high (red) or low (blue) A_SP_ in the space of location variance and the distance to the primary areas. This analysis reveals two *bands*: In low to medium areas there is a band of cortical regions exhibiting predominantly low A_SP_. At the same time a diagonal band from areas with low variance close to the primary areas, to higher variance, and intermediate distance to primary areas exhibits predominantly high A_SP_. Those areas include the middle temporal and the rostral middle frontal cortex on both hemispheres as well as parts of the superior frontal and inferior parietal cortex.

**Figure 4:**
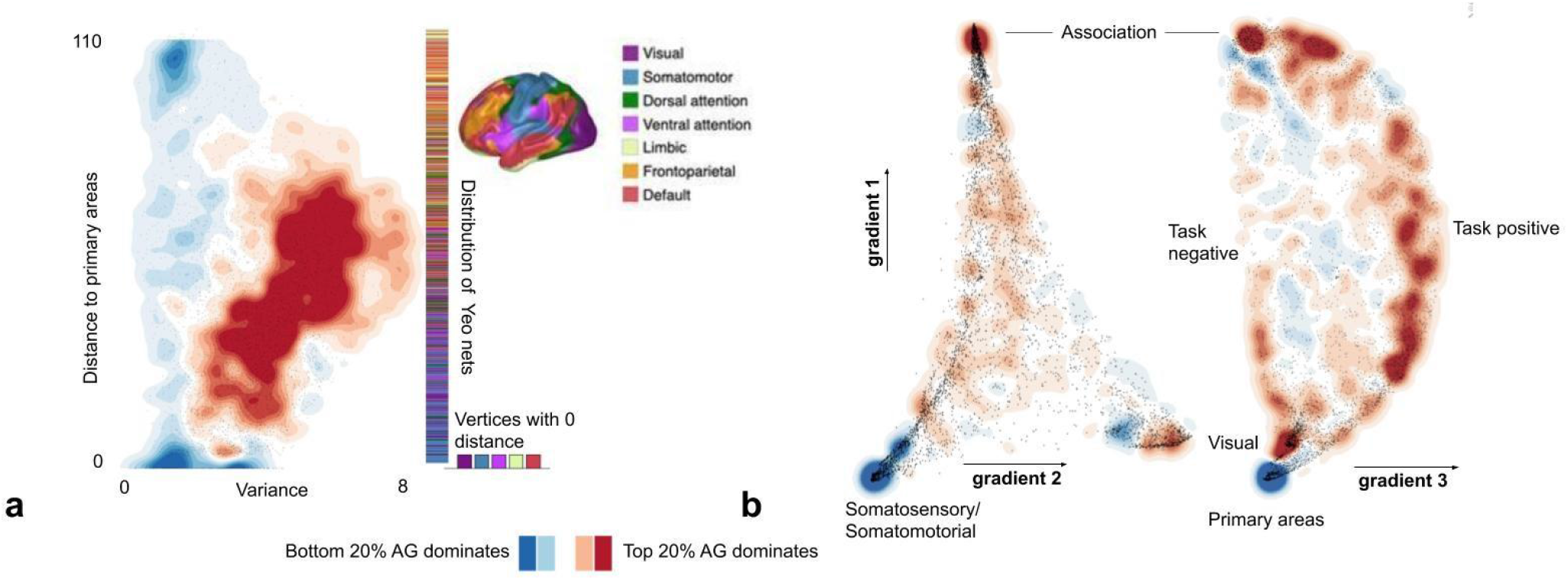
Relationship of spatial variability and functional gradients to genetic contribution to spatial layout. (a) Areas where high- (top 20%) and low- (bottom 20%) A_SP_ dominate are plotted in a coordinate frame spanned by the variance of the spatial position of network areas and the distance to primary areas. (b) Analogously, A_SP_ dominance maps are visualized in the functional gradient space for gradients 1, 2, and 3 (Margulies et al. 2016).

### 3.4 High genetic contribution to topography in areas in-between primary cortex and brain hubs

To further examine A_SP_ in the space of functional gradients spanning uni- to heteromodal, and task negative- to task positive networks, we plot high (red) or low (blue) A_SP_ dominance in the space spanned by the first, second and third functional gradients ^37^ (Figure 4b). In regions corresponding to association areas (high gradient 1), around visual processing (high gradient 2), and task positive activity (low gradient 3) high A_SP_ dominates locally, while in somatosensory and somatomotor areas (low gradient 1 and 2) low A_SP_ dominates. Task negative areas exhibit a mixed composition without a clear dominant behavior with regard to A_SP_. A *stream* of high (red) A_SP_ is situated along the edge from somatosensory/somatomotor to association areas through task positive regions. High genetic contributions modestly dominate the sparse central area of intermediate gradient 1 and 2.

### 3.5 Distinct strength and topography both contribute to overall variability

The overall variability as measured by the variance of aFC and FC estimated by the twin model is similar in magnitude (range aFC: 0.102 to 0.8, range FC: 0.105 to 0.76), and the genetic contributions A_aFC_ and A_FC_ moderately correlated on both hemispheres (left: r=0.298, corrected CI=(0.24,0.35); right: r=0.273, corrected CI=(0.21,0.32)). The correlation between A_aFC_ and A_SP_ was 0.128 on the left and 0.198 on the right hemisphere (corrected CI=(0.06,0.18) and corrected CI=(0.13,0.25)), suggesting that part of the variability of aFC and the corresponding additive genetic contribution is actually due to variability of the spatial layout of functional units on the cortex. Disentanglement of functional connectivity and topography reduces the correlation of genetic contributions to 0.05 (A_FC_ to A_SP_) on the left- and 0.091 on the right hemisphere (corrected CI=(−0.006,0.098) and CI=(0.03,0.15)). The variability explained by the twin model for SP ranges from 7.98 mm^2^ to 137 mm^2^ when described as variance. This corresponds to a standard deviation ranging from 2.82 mm to 11.7 mm. Supplementary Table 2 summarizes correlations and their confidence intervals. The hemispheres show similar patterns.

To gain a more detailed picture of the influence of FC and SP, we related their genetic contribution maps to cognitive functions, using a 12 component task activation model ^40^. (Figure 5). We were especially interested in comparing A_SP_ and A_FC_ in regions and corresponding functions, where both are high. Regions with high A_FC_ especially dominating over high A_SP_ are associated with visual processing and hand movements, whereas A_SP_ strongly dominates over A_FC_ in regions associated with inhibition and auditory processing.

**Figure 5:**
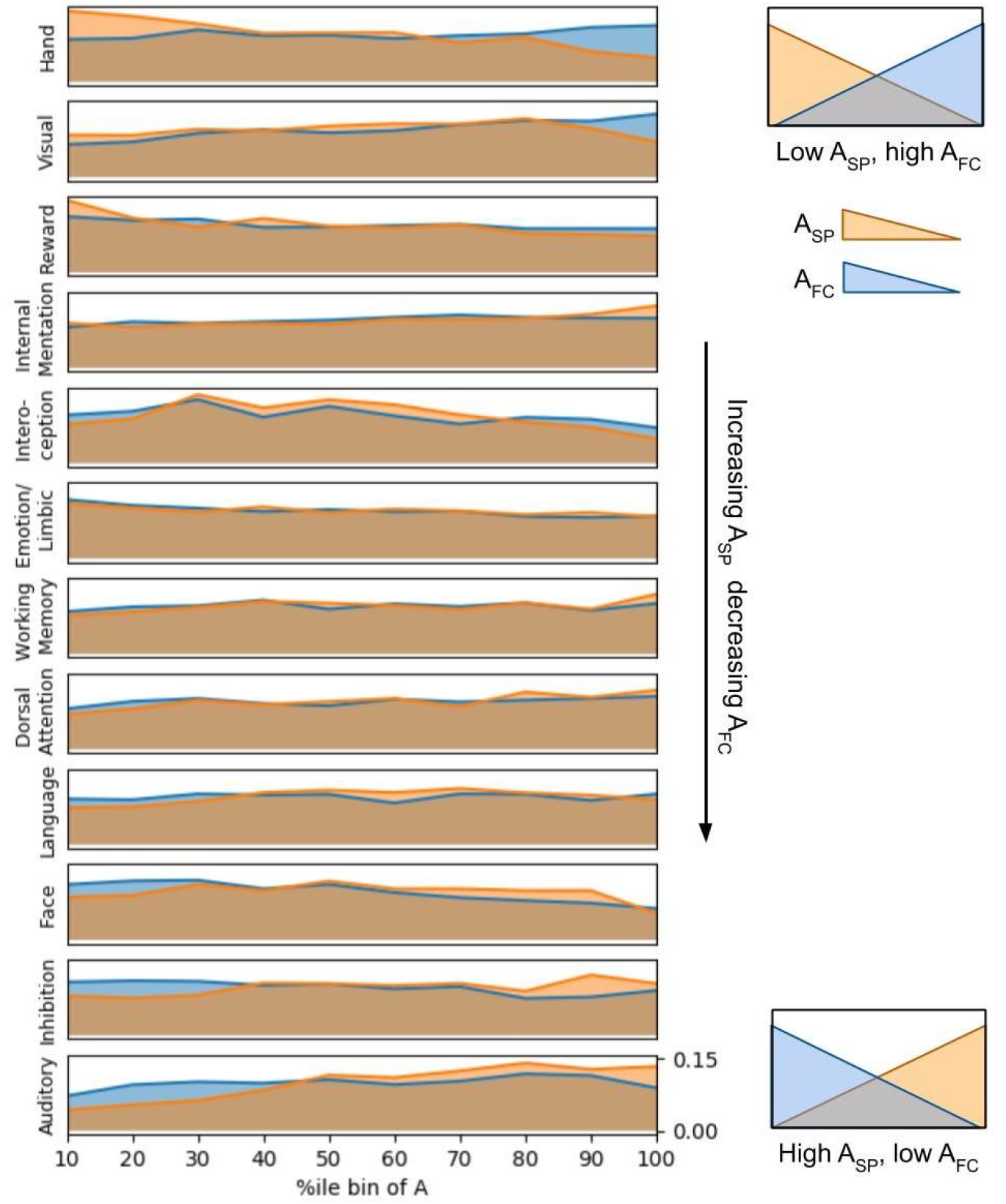
Relation of genetic contribution and cognitive function. Values of A_FC_ and A_PS_ are divided into 10 bins based on percentiles (x-axis). Within each bin normalized activation probabilities are summed given one of 12 components ^40^. The 12 components (rows) are ordered based on the difference of activation probabilities between A_FC_ and A_PS_ summed over bins and weighted by the log of percentile. The sequence visualizes the gradient from high A_SP_ and low A_FC_ on top, towards low A_SP_ and high A_FC_ on the bottom.

### 3.6 Replicability analysis

We performed replicability analysis using 2 RL fMRI runs, available for each participant in addition to the 2 LR runs used for the main discovery analysis. In contrast to the LR runs, the phase encoding direction for the RL runs is from right to left ^41^, which might be a source of variance between runs of the same subject, in addition to the general intra-subject variance ^8 42^. Disentangling resulted in comparable deformation fields of function and topography (Supplementary Figure 1). The variability of SP, FC and aFC estimated by the twin model at each vertex correlate in the discovery- and replicability analysis (Supplementary Figure 6). (aFC: left hemisphere: 0.98, right hemisphere: 0.98; FC: left: 0.93, right: 0.91; SP: left: 0.57, right: 0.65). The relationship between genetic and environmental influence, variability, and distance to primary areas, as well as their distribution in the functional gradient space is highly replicable (Supplementary Figure 7). Their respective dominance in 7 Yeo networks on the replication data, reflects those observed in the discovery data (Supplementary Figures 5).

## 4. Discussion

In this study we disentangled strength and topography of functional networks across the cortex. We applied a novel method to study the role of heritability and individual experience shaping these two key features independently. Disentanglement revealed that their influence on strength and topography diverges across the cortex. While in primary areas, strength is more heritable compared to topography, the opposite is the case in intermediate and higher order association areas. There, genetic factors primarily shape topography, while connectivity strength is predominantly determined by individual experience. Our results may inform our understanding of the mechanisms of emergence and continued adaptation of brain areas unique to humans.

### 4.1 Variable and heritable topography in intermediate and higher order association areas

In evolutionary older parts of the cortex, functional connectivity is related to proximity ^10^. The modern human association cortex does not follow this rule anymore. The *tethering hypothesis* ^11^ posits that regions that form modern association cortex became untethered from strong patterning signals of thalamic input in the past due to cortical expansion. This fostered a less hierarchical connectivity landscape in these regions. Together with an increase in cortico-cortical connectivity and their position between signal gradients from distributed primary areas, this may have paved the way for being subsequently co-opted for high-level integrative cognitive capabilities. In light of this, the rapid evolutionary expansion of association cortex may also have led to a dissociation between the roles of functional components and their spatial layout, contributing to the high variability these networks exhibit compared to primary areas ^8,10,11^. The precise role of this variability is, however, not clear.

We did observe dissociation of genetic influences on the variability of strength and topography in association cortex. While here, topography and connectivity strength are highly variable, the first is heritable to a much higher degree than the latter. Therefore the development of a less hierarchical connectivity landscape may have gone along with an increase in genetic variability encoding topographical variability. Since the genetic variability is still present today, different genotypes may have formed the basis of equally fit phenotypes in the past. At the same time, genetic influence may enable improving a coordinated - and eventually possibly even canonical - layout of processing pathways in the future, as heritability renders topographic variability visible to selection and adaptation, enabling the emergence of replicable organization. At the same time, the strength of the interconnectedness among the components anchored in a diverse, but heritable landscape is shaped by individual experience and the environment.

The observation that highly variable, and heritable topography overlaps in association cortex, is initially counter-intuitive, but may have benefits on the individual-, and the population level. It may afford connectivity strength’s variability and susceptibility to random environmental influence on top of an underlying coordinated framework of network topography, necessary for the acquisition of higher-order human cognitive abilities. At the same time, on the population level, simultaneous variability and heritability of topography may not only be a transient state while organization is optimising, but instead a means to sustain population level diversity and fitness^43^. For either point, the separation of variability explained by heritable traits versus random environmental influence and their independent link to topography and connectivity strength are important.

### 4.2 Coordinated processing pathways from primary to heteromodal areas

An increase of heritable topography from visual and somatomotor cortex to default and fronto-pariental networks (Figure 3c) supports the two hypotheses regarding different but coordinated heritable integration pathways. Task positive areas of high genetic contribution are in close proximity of early to intermediate processing areas found in a study of pathways from primary areas to network hubs ^44^. When taking into account areas influenced by common environment, high non-random contribution areas are next to the dorsolateral prefrontal cortex (DLPFC), the frontal eye fields (FEF) and a stream following the borders of the visual cortex to the lateral occipitotemporal junction (LOTJ) and then reaching into task-negative areas of the temporal lobe. Early integration areas of the visual processing stream form a cluster of high genetic contribution to SP present in the visual cortex in Figure 4b. The visual cortex’ connectivity follows three pathways ^44^, a feature not present for the other primary cortices. Genetic coding of network position in the visual cortex may be needed to robustly sustain a more intricate pathway structure. Low to intermediate integration areas exhibit comparably high heritability of functional connectivity strength (FC), though to a lesser extent. Peak areas are close to LOTJ and DLPFC, two regions belonging to integration areas not directly adjacent to the somatomotor and somatosensory cortex, and in case of the DLPFC not adjacent to the visual cortex ^44^. The stronger genetic encoding of connectivity strength in these areas might be necessary to effectively bridge the larger distance compared to areas located directly next to primary areas.

### 4.3 Disentanglement sheds new light on heritability of entangled functional connectivity strength

The entire cortex shows moderate genetic influence on connectivity strength before disentanglement (aFC). Disentangling FC and SP reveals a predominantly decreased FC compared to aFC and a location dependent increase of SP. This suggests that genetic influence on aFC to a relevant extent actually reflects A_SP_. Further evidence on that is provided in a recent study distinguishing connectivity profiles of sibling pairs from pairs of unrelated individuals ^45^. Although the study did not investigate heritability of topography directly, the authors used a parcellation to obtain regions of interest (ROIs) and described the time series of each ROI by the time series of all other ROIs through regression. They then used the obtained coefficients as a connectivity profile of a ROI. Most areas of low random environmental contribution to spatial topography overlap with cortical regions among the top 20 to distinguish sibling pairs from pairs of unrelated individuals ^45^ (Supplementary Figure 8).

This complements our emerging understanding of the role of variability of brain networks. As found in a recent study, variability in overall functional connectivity is actually reflected in the variability of topographical organisation and the spatial arrangement of functional regions strongly predicts behaviour ^46^. In another recent study ^47^ the entire cortical parcellation based on network topography was able to predict measures of cognition, personality and emotion. Together with our findings this suggests that different behavioral traits might be genetically determined through topography and have been either equally advantageous under similar conditions or alternatingly advantageous under changing conditions during human evolution.

Even though A_SP_ seems to determine A_aFC_ for some parts of the cortex, A_FC_ also has its role when looking at specific cognitive functions. Whereas genetic influence on most cognitive functions can be attributed to a mixture of A_SP_ and A_FC_ without a clear dominance of one of the two, there are also some exceptions as indicated by differences in ranking in Figure 5. Dorsal attention and language seem to be influenced genetically mainly through SP, since those functions are ranked at the bottom. Visual and hand movement related processing, on the other hand, are influenced genetically mainly through FC.

### 4.4 Related work

A range of papers has investigated genetic determination of morphology ^48, 49^ as well as structural ^16,17^ and functional features of the brain ^50, 15, 14, 45^. In a study of White et al. ^48^ cerebral brain volumes of twins, including whole brain volume as well as divisions into lobes and tissue types, showed correlations above 0.90, except the frontal white matter and occipital gray matter volume. Genetic correlations for surface area with different seed locations have been explored in ^49^ and showed an anterior-posterior division and a lack of long distance correlations. Similar to surface area, cortical thickness exhibited local genetic correlation with additional strong genetic correlation to homologs on the opposite hemisphere in another study ^51^. Glahn et al ^50^ explored heritability of anatomical morphology and functional aspects of the default mode network, revealing an independent genetic influence on anatomical morphology and functional variability. Some other previous studies based on fMRI data used a univariate twin model to investigate the connectivity of single edges between regions distributed across the cortex ^15, 14^. Observed genetic contributions are higher than common environmental influences in both papers. A multivariate machine learning based approach was used in ^45^ to distinguish sibling pairs from pairs of unrelated individuals. Useful regions for pair classification belonged to higher order systems, such as the fronto-parietal, dorsal attention and default mode network. Measures of structural connectivity were found to be especially heritable for connections within the default mode network, visual circuits and connections between default mode and fronto-parietal or ventral attention network ^16,17^.

### 4.5 Limitations

This study has several limitations. One limitation is the way how the genetic, common environmental and random environmental contributions to connectivity strength at each vertex are estimated. Since the brain is a network, describing the connectivity strength or profile of each vertex should preferably be done by including the connections to all nodes in the brain. However, due to the large number of parameters which had to be estimated for the twin model, this option is computationally infeasible. Therefore we describe the connectivity profile of each vertex by its connectivity to predefined regions of interest, an approach consistent with prior work ^31^. A limitation of the twin model used is that it only models contributions that act additively on the inter-subject variability. Including interactions of genetic and environmental influences in the model could give additional insights. Further limitations are that the power to separate A form C is lower than for separating A from E, which can lead to the partial attribution of common environmental influence to genetic influence or vice-versa. These concerns have been addressed through replicability analysis, which showed that the main findings are stable across runs and suggests that the processing stream from unimodal to heteromodal areas shows an interplay of rather genetically determined topography and functional connectivity strength determined rather by random environment, which deserves more investigation in the future.

## Abbreviations

aFC: variability of functional connectivity before disentanglement
SP: variability of topography of corresponding functional regions
FC: variability of functional connectivity strength of corresponding functional regions
A_aFC_, A_SP_, A_FC_: genetic contribution to indicated type of variability
E_aFC_, E_SP_, E_FC_: random environmental contribution to indicated type of variability

## Acknowledgements

We thank Oscar Miranda-Dominguez for sharing his version of the Gordan parcelation with us. This work was supported by the Medical University of Vienna.

