## Supplementary material for "Disentangling cortical functional connectivity strength and topography reveals divergent roles of genes and environment"

#### 1 Replicability Analysis

We performed replicability analysis using 2 RL fMRI runs, which have also been available for each participant additionally to the 2 LR runs used for the discovery analysis. In contrast to the LR runs, the phase encoding direction for the RL runs is from right to left [27]. First, we compared the variability of SP, FC and aFC estimated by the twin model at each vertex in the discovery and replicability analysis (Maps on discovery- and replicability data are shown in Supplementary Figure 6). For aFC, the correlation of the cortical variability maps was around 0.98 on both hemispheres. The correlation is slightly lower for FC with 0.93 on the left and 0.91 on the right hemisphere. Variability maps for SP are the least similar (left: 0.57, right:0.65). While on the vertex-level maps for A, C and E exhibit less correlation, in concordance with the discovery analysis, SP shows the broadest range of genetic influence across the cortex (0% to 62.8%), with peak values higher than for aFC and FC, whose range of genetic contribution values is 4.66% - 46.8% and 4.2% - 39.7%, respectively. Again, the average genetic contribution to FC calculated over all vertices on the cortex decreased significantly (left hem: 19.5% (aFC) vs 16.5 % (FC) of variance explained by genes, corrected p-value <0.0001; right hem: 19.3% (aFC) vs 16.5% (FC) of variance explained by genes, corrected p-value < 0.0001). Furthermore, random environment is still the dominating influence on the cortex for FC, whereas SP shows higher genetic contribution than random environment at peaks.

Vertex-wise genetic (A), common environmental (C) and random environmental (E) contributions to variability of aFC, FC, and SP are shown for both hemispheres in Supplementary Figures 3 (discovery) and 9 (replicability analysis). A, C and E - vertex maps for aFC correlate in a range from 0.362 (A, left hemisphere) - 0.631 (E, right hemisphere), vertex maps for FC correlate in a range from 0.153 (C, left hemisphere) to 0.354 (E, right hemisphere), and vertex maps for SP correlated with values from 0.021 (A, left hemisphere) to 0.405 (E, right hemisphere).

For the replication analysis on the RL runs of each individual plots similar to Figure 4 (Supplementary Figure 7) for both hemispheres are given in the supplementary material. The positive correlation with variability of network position is the same (Supplementary Figure 7a; left hemi: $r = 0.2$, corrected $p < 0.0001$; right hemi: $r=0.33$, corrected $p< 0.0001$) as well as the presence of a stream of high genetic contribution along the edge of association areas leading to the task positive related top of the three-sided pyramid formed by component 1-3 Supplementary Figure 7b). The same holds for the cluster of high genetic contribution in the corner representing the visual areas. Furthermore, for both analyses areas where genetic influence to SP especially dominates, belong to the fronto-parietal the attention and the default mode network (Supplementary Figure 5).

Although some patterns of genetic influence on SP are stable, the cortical maps of variability and its split in A, C and E are less similar than those for FC and aFC. A reason for the instability of cortical maps may lie in the difference of the phase encoding direction between LR (left-right) and RL (right-left) resting state fMRI data, which might be a source of variance between runs of the same subject, in addition to the general intra-subject variance [8, 28].

### 2 Supplementary Figures

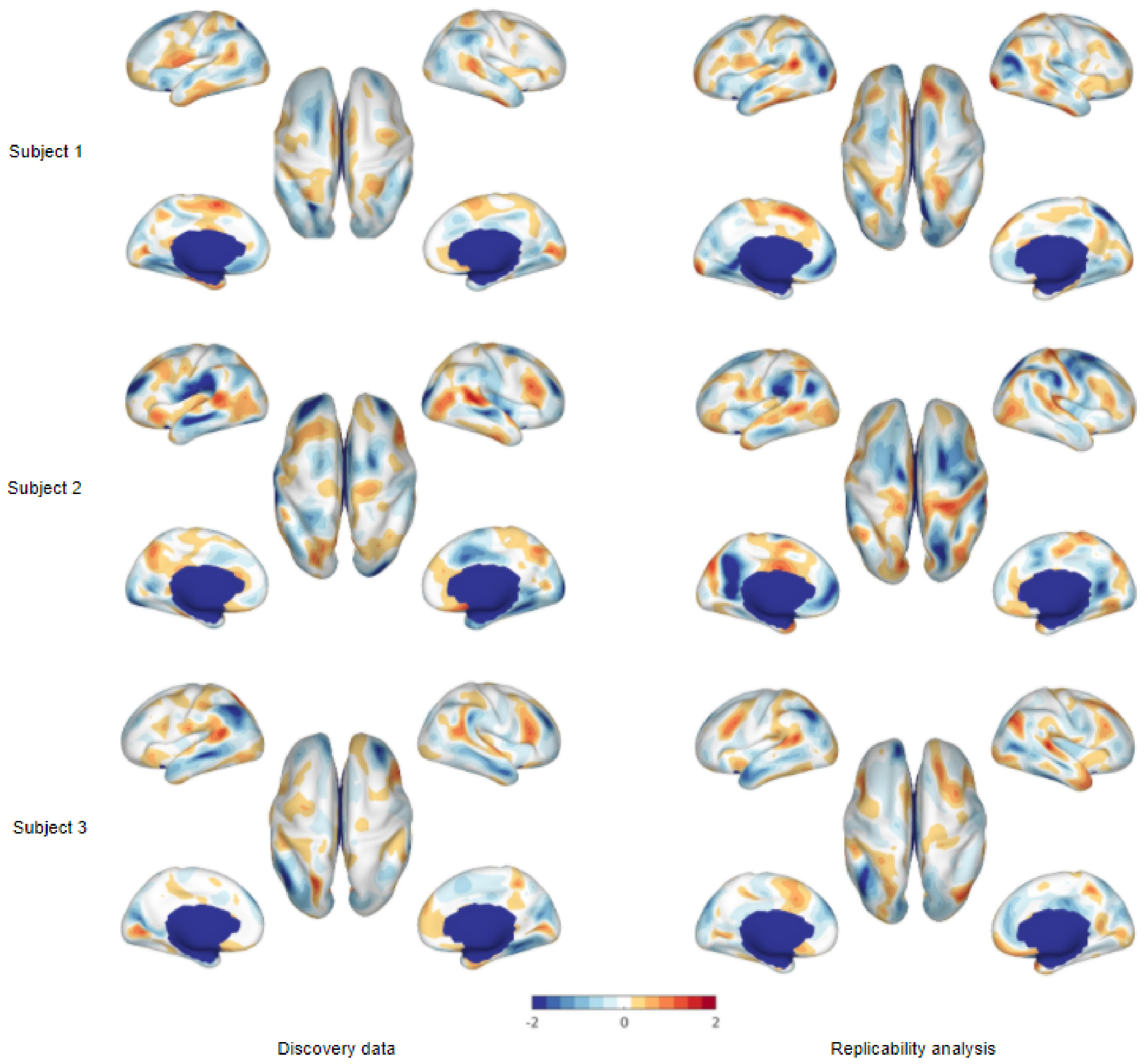


Supplementary Figure 1: Deformation fields of exemplary subjects after functional alignment. Shown is the areal distortion for each vertex given as weighted average of face distortions of this vertex averaged across subjects. Rows correspond to subjects columns to discovery data (left) and replicability analysis (right)

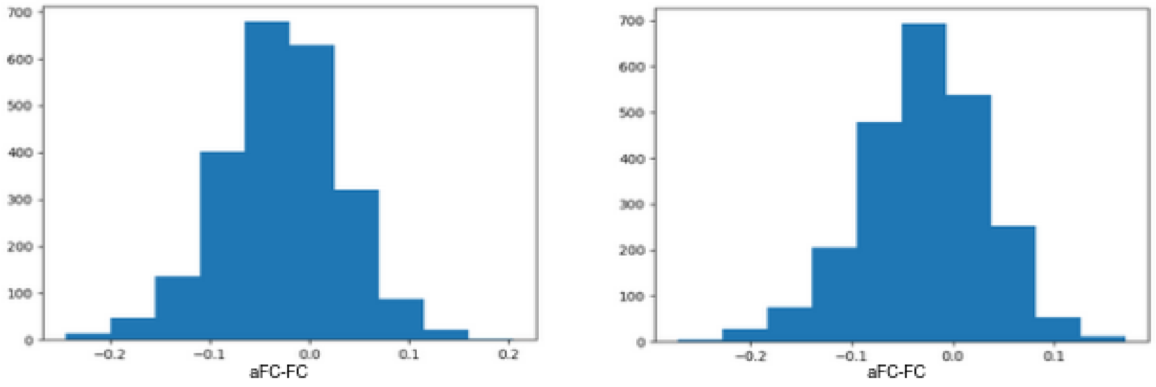


Left hemisphere

Right hemisphere

Supplementary Figure 2: Histograms showing the distribution of aFC-FC to check the normality assumption of the paired t-test. On both hemispheres the differences are normally distributed.

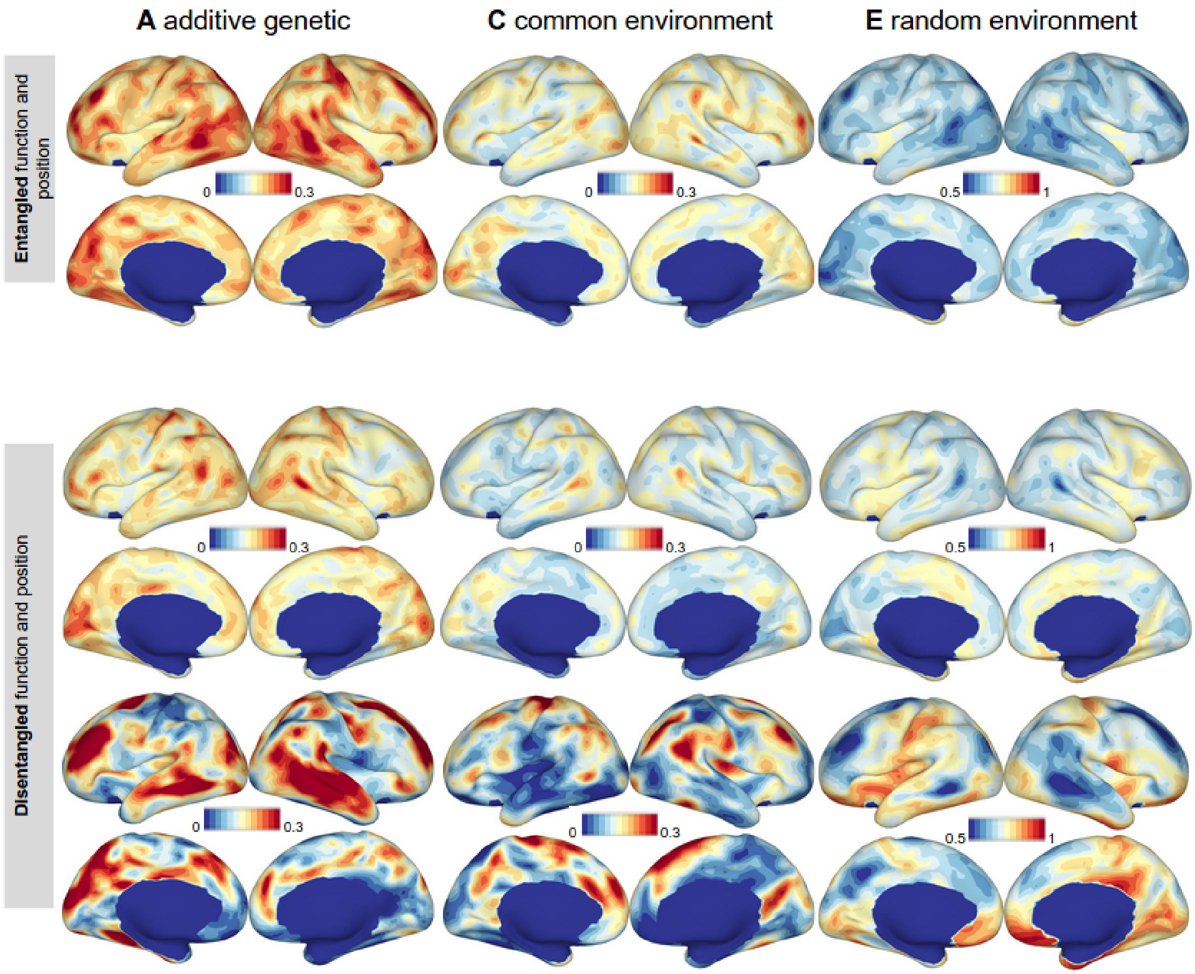


<u>Supplementary Figure 3:</u> Additive genetic, common environmental and random environmental contribution for entangled and disentangled function and topography. The contribution is presented as fraction of the total variance explained by the twin model. The color scale is the same within each column. Since high genetic contribution to SP is attributed only to some spots, we want to take a closer look at the landscape of contributions for SP. The peak estimates are higher or equal to 34.0% for additive genetic and higher or equal to 27.1% for common environmental contribution of the total variance explained. The areas with peak estimates on the left hemisphere for non random contribution are the rostral middle frontal region, the middle temporal region, the inferior parietal region and the superior part of the precentral region on the lateral brain and the precuneus as well as a spot in the superior frontal region on the medial side. On the right hemisphere again the middle temporal region and a part of the rostral middle frontal region exhibit high values. However, the boundary region between superior frontal and caudal middle frontal area as well as the boundary area between inferior parietal and supa marginal region exhibit high genetic contribution to SP on the lateral right hemisphere. On the medial side there are similar but smaller clusters as on the left hemisphere. On both hemispheres the somatomotor area and transverse temporal area on the lateral side as well as the medial orbital frontal region, the para central lobule, the lingual and pericalcarine region on the medial side show high contributions of random environment. On the right hemisphere the posterior cingulate also shows high random contributions which are present in part of the isthmus and precuneus, too.

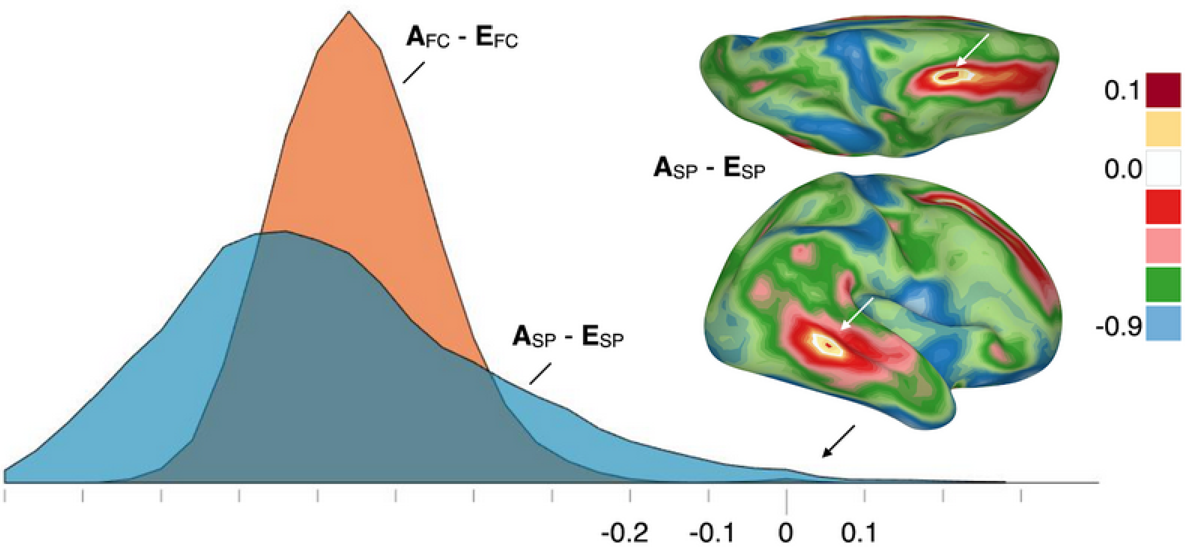


Supplementary Figure 4: Comparison of genetic and random environmental influence at each cortical location. Two hotspots where genetic influence on SP is higher than random environmental influence are revealed.

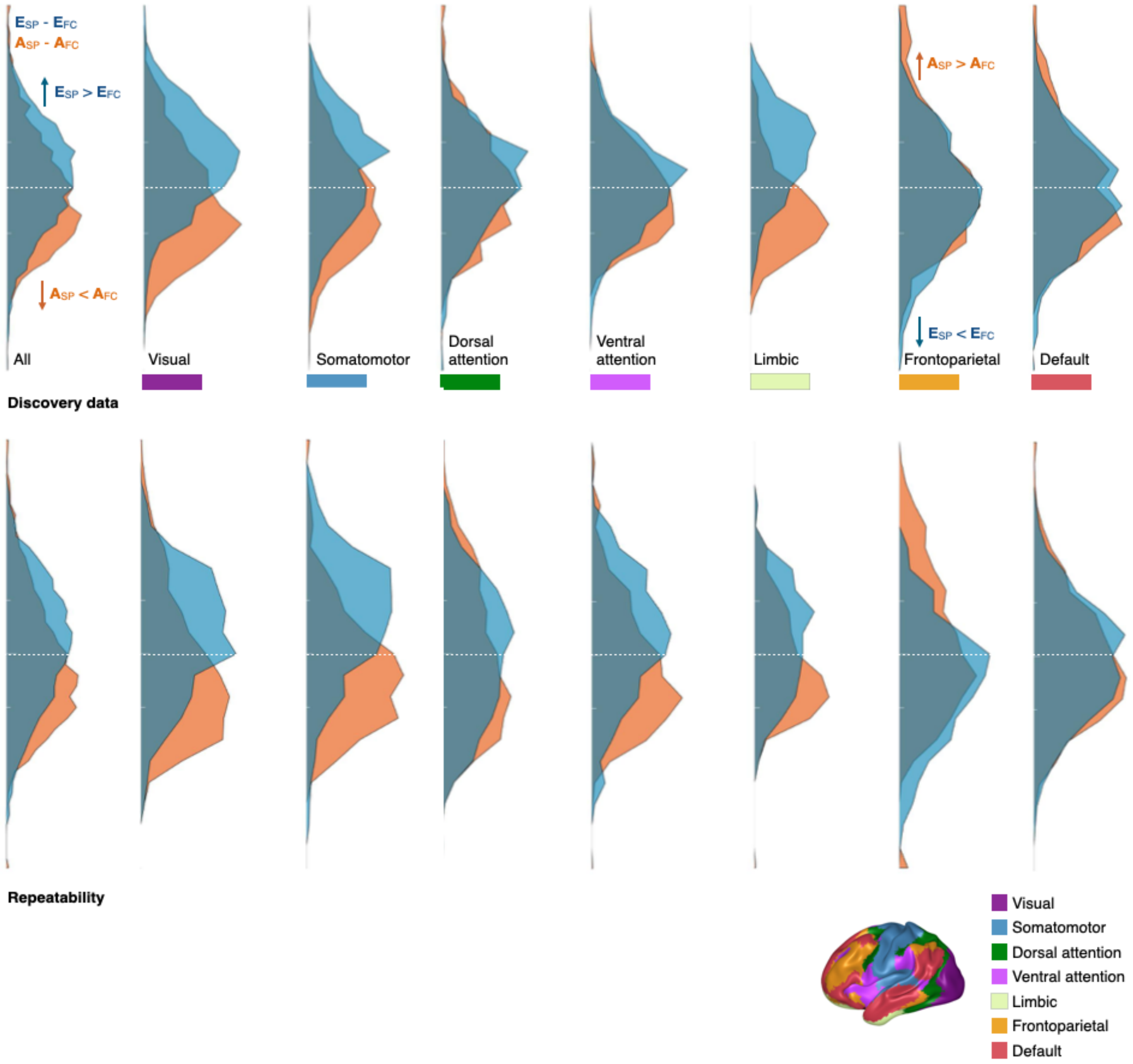


Supplementary Figure 5: Distribution of $A_{SP}$-$A_{FC}$ and $E_{SP}$-$E_{FC}$ for various networks for discovery data and replicability analysis. For both analyses, spots where $A_{SP}$ especially dominates over $A_{FC}$ belong to the attention networks the fronto-parietal and the default mode network. In the visual and somatomotor cortex mainly $A_{FC}$ dominates.

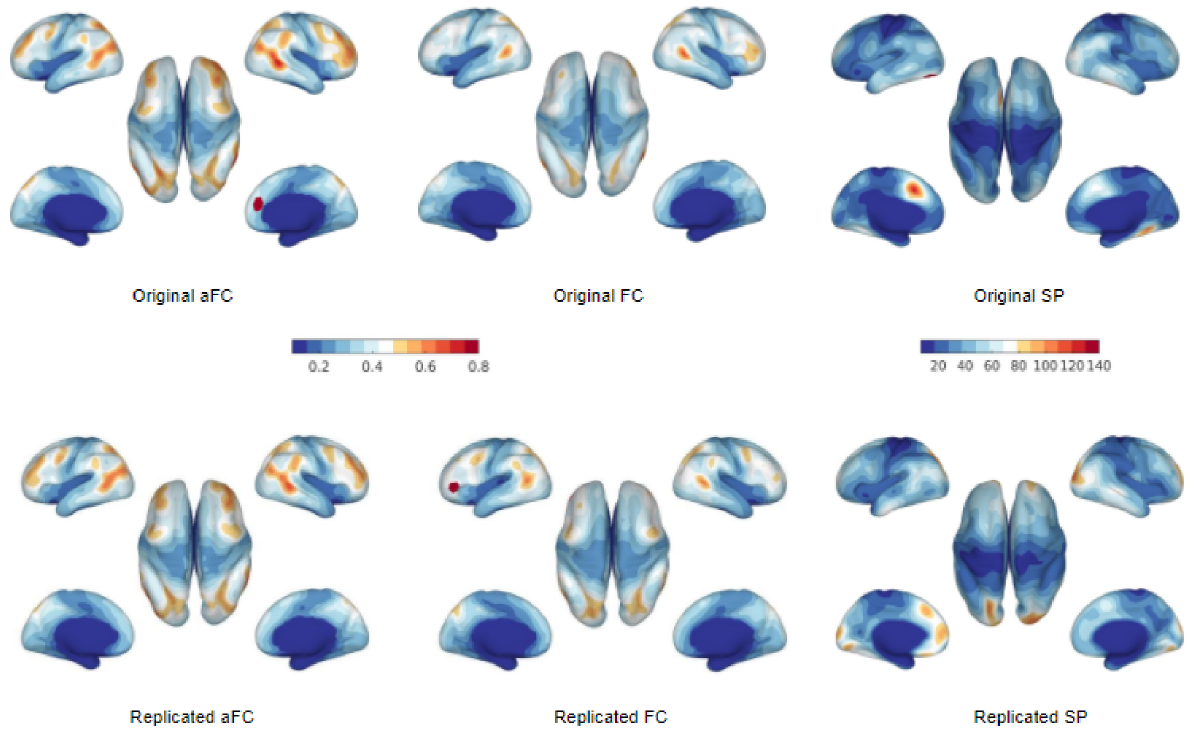


Supplementary Figure 6: Variability estimated by twin model for aFC, FC and SP. The top row shows results for the discovery analysis, the bottom row shows the results for the replicability analysis.

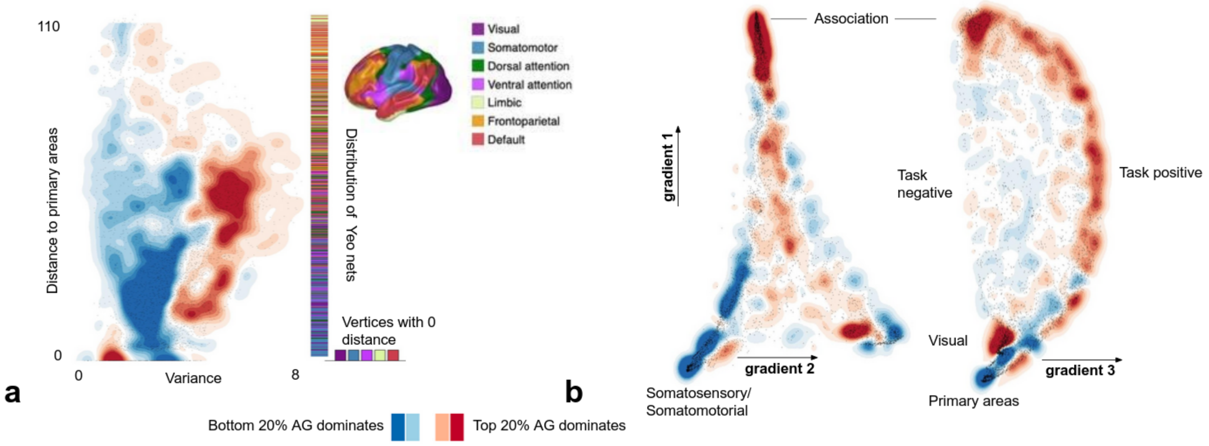


Supplementary Figure 7: Pendant to Figure 4 for the replicability analysis. (a) SP variance and distance to primary areas of cortical regions where points with high (top 20%) and low (bottom 20%) $A_{SP}$ dominate; (b) Analogous $A_{SP}$ dominance maps for gradients 1, 2, and 3.

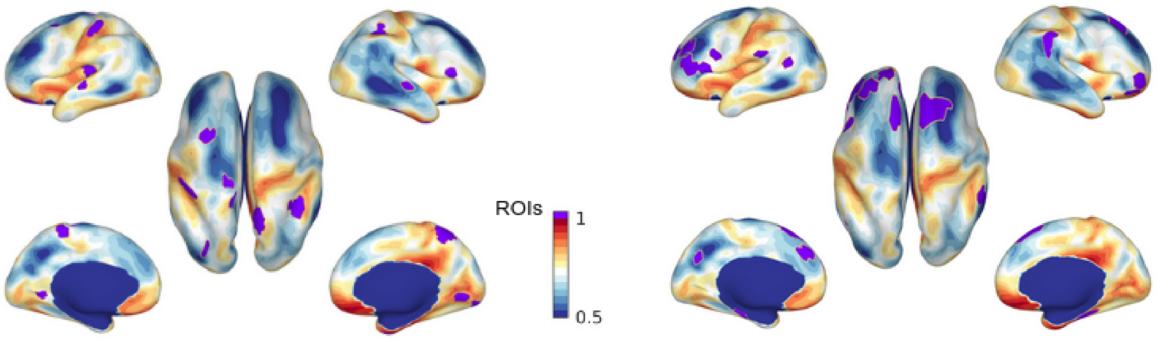


ROIs ranked under top 20 in Miranda-Dominguez et al. 2018

ROIs ranked under least 20 in Miranda-Dominguez et al. 2018

Supplementary Figure 8: ROIs belonging to Gordon atlas overlayed on random environmental influence to SP. On the left, ROIs ranked under top 20 in Miranda-Diminguez et al. 2018 to distinguish sibling pairs from pairs of unrelated individuals are plotted. The ROIs mainly overlap with areas of low (blue) random environmental influence. On the right handside the lowest ranked 20 cortical ROIs are plotted. They overlap mainly with areas of high (orange) random environmental contribution.

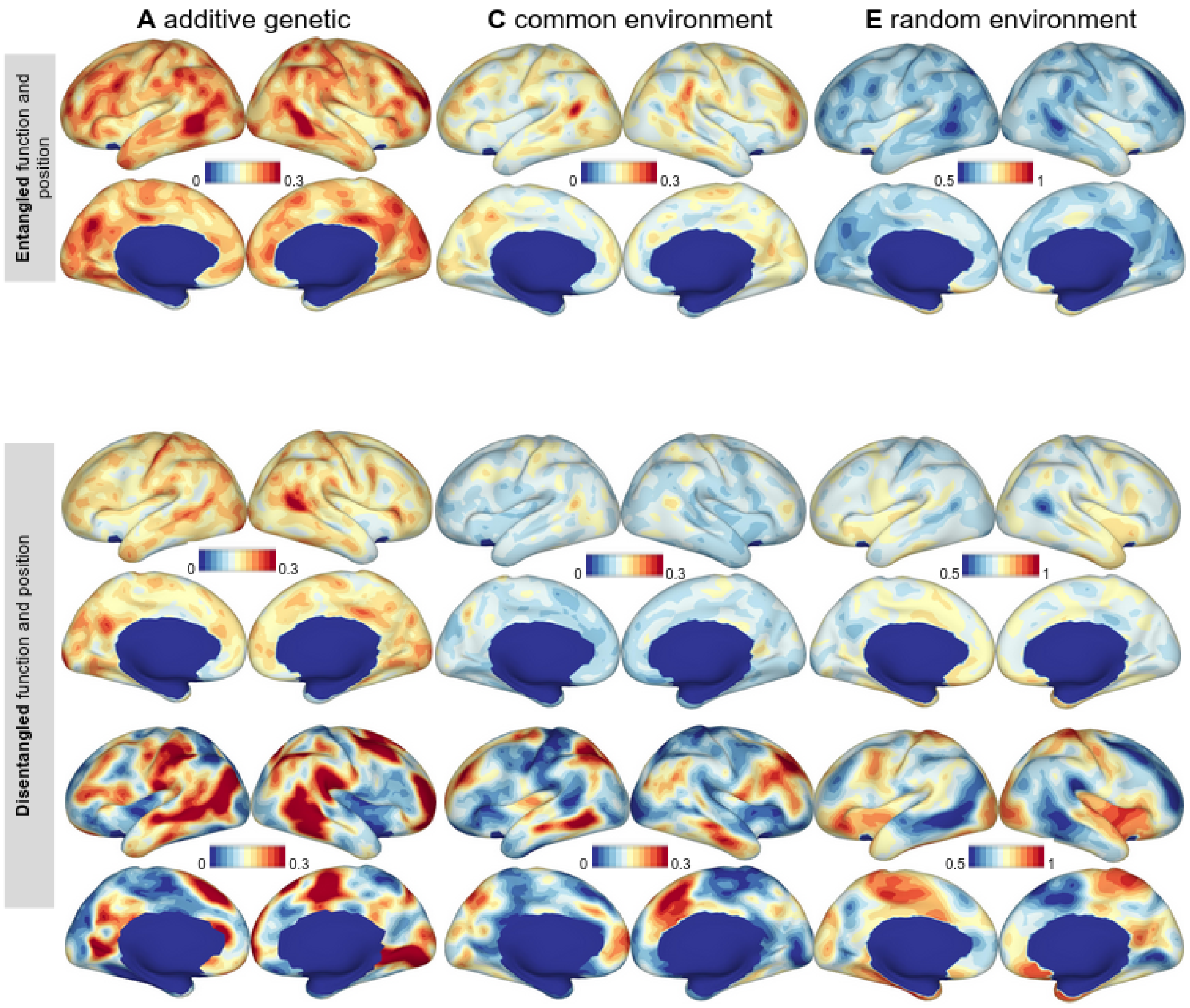


Supplementary Figure 9: Additive genetic, common environmental and random environmental contribution for entangled and disentangled function and topography - replicability analysis. The contribution is presented as a fraction of the total variance explained by the twin model. The color scale is the same within each column.

### 3 Supplementary tables

**Supplementary Table 1:** Correlations of genetic contribution map with various cortical surface maps

| Map to correlate | Correlation value | Uncorrected confidence interval | Corrected confidence interval |
|---|---|---|---|
| **Left hemisphere** | | | |
| Variability of spatial layout | 0.30 | (0.26, 0.34) | (0.24,0.35) |
| Distance to primary area (somatomotor/-sensory, visual, ventral and dorsal attention) | 0.21 | (0.15, 0.0.26) | (0.13,0.27) |
| Distance to primary area (limbic, fronto-parietal, default) | -0.27 | (-032,-0.23) | (-0.35,-0.22) |
| Cortex expansion | 0.18 | (0.13, 0.23) | (0.11,0.24) |
| **Right Hemisphere** | | | |
| Variability of spatial layout | 0.41 | (0.38, 0.45) | (0.36,0.46) |
| Distance to primary area (somatomotor/-sensory, visual, ventral and dorsal attention) | 0.29 | (0.23, 0.34) | (0.21,0.35) |
| Distance to primary area (limbic, fronto-parietal, default) | -0.01 | (-0.06, 0.05) | (-0.09, 0.06) |
| Cortex expansion | 0.099 | (0.06, 0.14) | (0.03,0.15) |

**Supplementary Table 2:** Correlations of genetic contribution maps:

| Map to correlate | Correlation value | Uncorrected confidence interval | Uncorrected confidence interval |
|---|---|---|---|
| **Left hemisphere** | | | |
| Strength (anat.)/ strength (func.) | 0.298 | (0.26, 0.34) | (0.24,0.35) |
| Strength (anat.)/ position | 0.128 | (0.09, 0.17) | (0.06,0.18) |
| Strength (func.)/ position | 0.05 | (0.009, 0.09) | (-0.006,0.098) |
| **Right hemisphere** | | | |
| strength (anat.)/ strength (func.) | 0.273 | (0.23, 0.31) | (0.21,0.32) |
| strength (anat.)/ position | 0.198 | (0.16, 0.24) | (0.13,0.25) |
| strength (func.)/ position | 0.091 | (0.05, 0.13) | (0.03,0.15) |
